# Beyond antibiotic resistance: the *whiB7* transcription factor coordinates an adaptive response to alanine starvation in mycobacteria

**DOI:** 10.1101/2023.06.02.543512

**Authors:** Nicholas C. Poulton, Michael A. DeJesus, Vanisha Munsamy-Govender, Cameron G. Roberts, Zachary A. Azadian, Barbara Bosch, Karl Matthew Lin, Shuqi Li, Jeremy M. Rock

## Abstract

Pathogenic mycobacteria are a significant cause of morbidity and mortality worldwide. These bacteria are highly intrinsically drug resistant, making infections challenging to treat. The conserved *whiB7* stress response is a key contributor to mycobacterial intrinsic drug resistance. Although we have a comprehensive structural and biochemical understanding of WhiB7, the complex set of signals that activate *whiB7* expression remain less clear. It is believed that *whiB7* expression is triggered by translational stalling in an upstream open reading frame (uORF) within the *whiB7* 5’ leader, leading to antitermination and transcription into the downstream *whiB7* ORF. To define the signals that activate *whiB7*, we employed a genome-wide CRISPRi epistasis screen and identified a diverse set of 150 mycobacterial genes whose inhibition results in constitutive *whiB7* activation. Many of these genes encode amino acid biosynthetic enzymes, tRNAs, and tRNA synthetases, consistent with the proposed mechanism for *whiB7* activation by translational stalling in the uORF. We show that the ability of the *whiB7* 5’ regulatory region to sense amino acid starvation is determined by the coding sequence of the uORF. The uORF shows considerable sequence variation among different mycobacterial species, but it is universally and specifically enriched for alanine. Providing a potential rationalization for this enrichment, we find that while deprivation of many amino acids can activate *whiB7* expression, *whiB7* specifically coordinates an adaptive response to alanine starvation by engaging in a feedback loop with the alanine biosynthetic enzyme, *aspC*. Our results provide a holistic understanding of the biological pathways that influence *whiB7* activation and reveal an extended role for the *whiB7* pathway in mycobacterial physiology, beyond its canonical function in antibiotic resistance. These results have important implications for the design of combination drug treatments to avoid *whiB7* activation, as well as help explain the conservation of this stress response across a wide range of pathogenic and environmental mycobacteria.

## INTRODUCTION

Mycobacteria cause a wide range of diseases and include some of the oldest infections described in human history, leprosy and tuberculosis (TB). Both of these diseases are still prevalent today^1, 2^, and TB remains the leading cause of death from any single infectious agent^2^. Infections caused by non-tuberculous mycobacteria, including *Mycobacterium abscessus* and *Mycobacterium avium* are increasingly common, especially amongst immunocompromised individuals^3^. Although these mycobacterial diseases vary in manifestation, they share the common feature of being difficult to treat.

The reasons why mycobacterial diseases are difficult to treat are complex and multifactorial^4–7^. However, one major difficulty is the high level of intrinsic drug resistance of mycobacteria^8^. Intrinsic drug resistance can be attributed in part to the relatively impermeable mycobacterial outer membrane, which is composed of a thick layer of waxy mycolic acids and other glycolipids which slow the uptake of a wide range of compounds^9, 10^. Further, mycobacteria have a large number of additional intrinsic drug resistance mechanisms that act via drug efflux^11^, drug modification^12^, target modification^13^, and target rescue^14^. Although intrinsic drug resistance determinants vary across mycobacterial species, the *whiB7* pathway has emerged as a conserved and key mechanism limiting the activity of different antibiotics against diverse mycobacterial pathogens^15–18^. WhiB7 is a transcriptional activator that promotes the expression of a suite of intrinsic drug resistance determinants including: 1) the *tap* drug efflux pump^19^; 2) *eis*, an acetyltransferase that modifies and inactivates aminoglycosides^12^; 3) *erm*, a ribosomal RNA methyltransferase whose activity prevents macrolide antibiotic engagement with the ribosome^13^; 4) *hflX*, a ribosome splitting factor that rescues drug-stalled ribosomes^14^; and many other genes^17^.

WhiB7 was originally discovered in the soil bacterium *Streptomyces lividans*^16^. *whiB7* was hypothesized to have originated as a self-protection mechanism in antibiotic-producing bacteria, and then been acquired and retained by an ancestral soil-dwelling actinobacteria to protect against antibiotics being produced by other soil bacteria^16^. Why WhiB7 would be retained in pathogenic mycobacteria like Mtb and *M. leprae*, long after their progenitors left the antibiotic containing soil, has remained an open question.

WhiB7 is activated in response to and protects against a number of different antibiotics, particularly ribosome-targeting antibiotics like macrolides, lincosamides, and spectinomycin^16^. *whiB7* expression is controlled by an upstream open reading frame (uORF)-mediated transcriptional attenuation mechanism in its 5’ regulatory region^20–22^. Under non-stressed conditions, *whiB7* transcription initiates at a distal upstream transcriptional start site (TSS) and proceeds through a short uORF. Efficient translation of the uORF during un-stressed conditions results in the formation of a rho-independent terminator which stops transcription prior to the start of the *whiB7* ORF^22^. During conditions of translation stress, for example the presence of a macrolide antibiotics, the uORF is inefficiently translated. This results in the formation of an antiterminator structure and transcriptional readthrough into the *whiB7* ORF^22^. Elevated WhiB7 levels then initiate a positive feedback loop by binding to and activating transcription from the *whiB7* promoter, generating *whiB7* mRNA levels as much as 1,000 times higher than basal *whiB7* expression under unstressed conditions^23, 24^.

Beyond ribosome-targeting antibiotics, numerous groups have identified epistatic mutations that result in constitutive *whiB7* activation and decreased drug sensitivity. Gomez et al. identified *M. smegmatis* mutants harboring mutations in *rplO*, *rplY*, and *rplF* that showed broad, low-level antibiotic resistance, in part due to constitutive *whiB7* activation^25^. Schraeder et al. identified *M. smegmatis* mutants in the arginine biosynthetic genes *argA* and *argD* that led to constitutive *whiB7* activation and promoted tolerance to aminoglycoside antibiotics^26^. Our group identified partial loss-of-function mutations in the essential translation factor *ettA* in Mtb clinical isolates that confer low-level resistance to multiple antibiotics, in part through constitutive *whiB7* activation^27^. Lastly, through yet to be defined mechanisms, WhiB7 is activated in response to host-derived stressors during *ex vivo* infection in macrophages^28^. Collectively, these findings suggest that alterations in various biological processes can lead to *whiB7* activation^23^.

Here, we set out to comprehensively define the biological pathways whose inhibition leads to *whiB7* activation. By coupling a *whiB7* expression reporter with a genome-wide CRISPRi screen, we identified ∼150 genes whose inhibition results in constitutive *whiB7* activation. In addition to the expected functions in ribosome biogenesis and translation, we identified a diverse set of hit genes involved in other cellular pathways including redox homeostasis and transcription. We show that the responsiveness of *whiB7* to amino acid starvation is determined by the amino acid composition of the uORF, which explains species-specific differences in *whiB7* activation signals. To identify physiological pathways beyond intrinsic resistance that may be dependent on *whiB7*, we performed a CRISPRi vulnerability screen. This screen identified a regulatory feedback loop where *whiB7* can sense the deprivation of alanine to upregulate expression of the alanine aminotransferase, *aspC,* demonstrating a physiological role for *whiB7* outside of its canonical role in intrinsic drug resistance.

## RESULTS

### Development of *whiB7* reporter systems in mycobacteria

To better understand the complex set of signals that can activate *whiB7*, we sought to globally identify genes whose inhibition results in constitutive *whiB7* activation. To identify such genes, we built two *whiB7* transcriptional reporters^23^. The entire *whiB7* regulatory region (500 bp upstream of the *whiB7* ORF start codon) was cloned upstream of either a zeocin resistance gene (*zeoR*) or the *mScarlet* fluorescent reporter (**Fig. 1A**). Consistent with faithful recapitulation of *whiB7* regulation, both reporters displayed the expected increase in activity upon inhibition of translation (**Fig. 1B-E**). Translation was inhibited either chemically with clarithromycin or genetically with knockdown of the essential translation factor *ettA* or the arginine biosynthesis gene *argD*.

**Figure 1.**
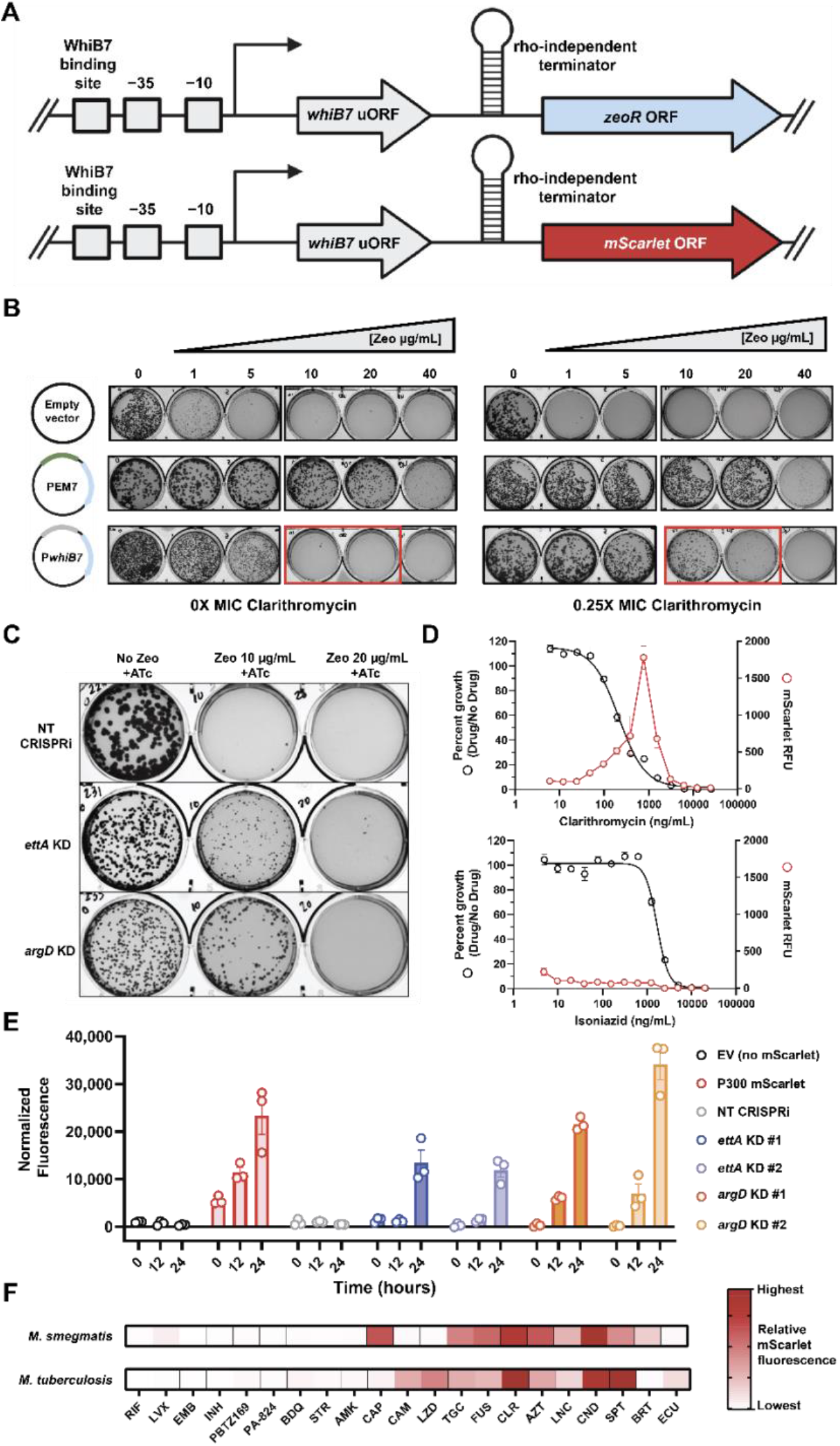
Development and validation of *whiB7* reporters. (A) Genetic architecture of *PwhiB7* reporter constructs. Both reporters were cloned into single-copy, integrating plasmids. (B) Zeocin resistance profiles of *M. smegmatis* strains (∼1,000 colony forming units (CFU)/well) harboring the zeocin resistance gene driven by the indicated promoters. EM7 is a strong, constitutive promoter. Plates contain either no drug (left) or 100 ng/ml = 0.25X minimum inhibitory concentration (MIC) of the known *whiB7* activating drug, clarithromycin (right). The red rectangle marks clarithromycin-dependent growth of the *M. smegmatis PwhiB7:zeoR* strain in the presence of 10-20 μg/ml zeocin. (C) Growth of the indicated *M. smegmatis* CRISPRi strains on agar plates containing zeocin at 0, 10, or 20 ug/mL. NT = non-targeting; KD = knockdown. (D) Dose response curves (mean ± s.e.m., n = 3 replicates) of the *PwhiB7:mScarlet* reporter *M. smegmatis* strain for clarithromycin (top graph) and isoniazid (bottom graph). Drug dose-response curves (percent growth) are shown in black and mScarlet fluorescence (RFU) are shown in red. (E) Normalized fluorescence values of the indicated *M. smegmatis* CRISPRi strains at 0, 12, and 24 hours after addition of ATc to activate CRISPRi (mean ± s.e.m., n = 3 replicates). Both *ettA* and *argD* are targeted with two different sgRNAs each, denoted #1 and #2. P300 is a strong, constitutive promoter. EV = empty vector. (F) Normalized fluorescence values for the *PwhiB7:mScarlet* reporter strain (top row: *M. smegmatis*, bottom row: *M. tuberculosis*) in response to the listed panel of drugs. Fluorescence values are indicative of the highest value obtained at a sub-MIC concentration of each drug. Normalized RFU: *M. smegmatis* = -18.2 to 2,000; *M. tuberculosis* = 13.2 to ≥6,000. RIF = rifampicin; LVX = levofloxacin; EMB = ethambutol; INH = isoniazid; BDQ = bedaquiline; STR = streptomycin; AMK = amikacin; CAP = capreomycin; CAM = chloramphenicol; LZD = linezolid; TGC = tigecycline; FUS = fusidic acid; CLR = clarithromycin; AZT = azithromycin; LNC = lincomycin; CND = clindamycin; SPT = spectinomycin; BRT = bortezomib; ECU = ecumicin^29^.

To further validate the sensitivity and specificity of the *mScarlet* reporter system, a larger panel of antibiotics were tested for their ability to induce *whiB7* in both *M. smegmatis* and *M. tuberculosis*. Results were largely concordant between the two bacterial species, with several ribosome-targeting drugs including macrolides, lincosamides, and spectinamides resulting in potent *whiB7* induction compared to antibiotics targeting other biological processes (**Fig. 1F, Fig. S1)**. For reasons yet unknown, *M. tuberculosis* and *M. smegmatis* showed subtle differences in *whiB7* induction by several drugs, including chloramphenicol, linezolid, and capreomycin, which may reflect species-specific differences in ribosome protection factors.

### Genome-scale CRISPRi epistasis screen identifies a diverse set of genes whose inhibition activates *whiB7*

To identify genes whose inhibition activates *whiB7*, we transformed an *M. smegmatis* strain harboring the *PwhiB7:zeoR* reporter construct with a genome-scale CRISPRi library (**Fig. 2A**). This library contains 176,451 sgRNAs targeting 6,642/6,679 of all *M. smegmatis* genes and includes sgRNAs of varying predicted knockdown efficiencies to enable tunable knockdown of essential genes^30^. The library was grown for ∼5 generations in the presence of anhydrotetracycline (ATc) to activate CRISPRi, induce target knockdown, and allow for *whiB7* activation and production of the ZeoR selectable marker. After this “pre-depletion” phase, the library was plated in the presence of ATc with or without zeocin. After outgrowth, roughly 100-fold fewer colonies were observed on the zeocin-containing vs lacking plates, suggesting that only ∼1% of all sgRNAs selected for *whiB7* expression (**Fig. 2B,C**). Spontaneous zeocin resistance, i.e. zeocin resistance in the absence of ATc, was a negligible contributor to the overall number of colonies (**Fig. 2B)**. Genomic DNA from colony forming units (CFU) from each set of plates was extracted and used to quantify sgRNA abundance by deep sequencing.

**Figure 2.**
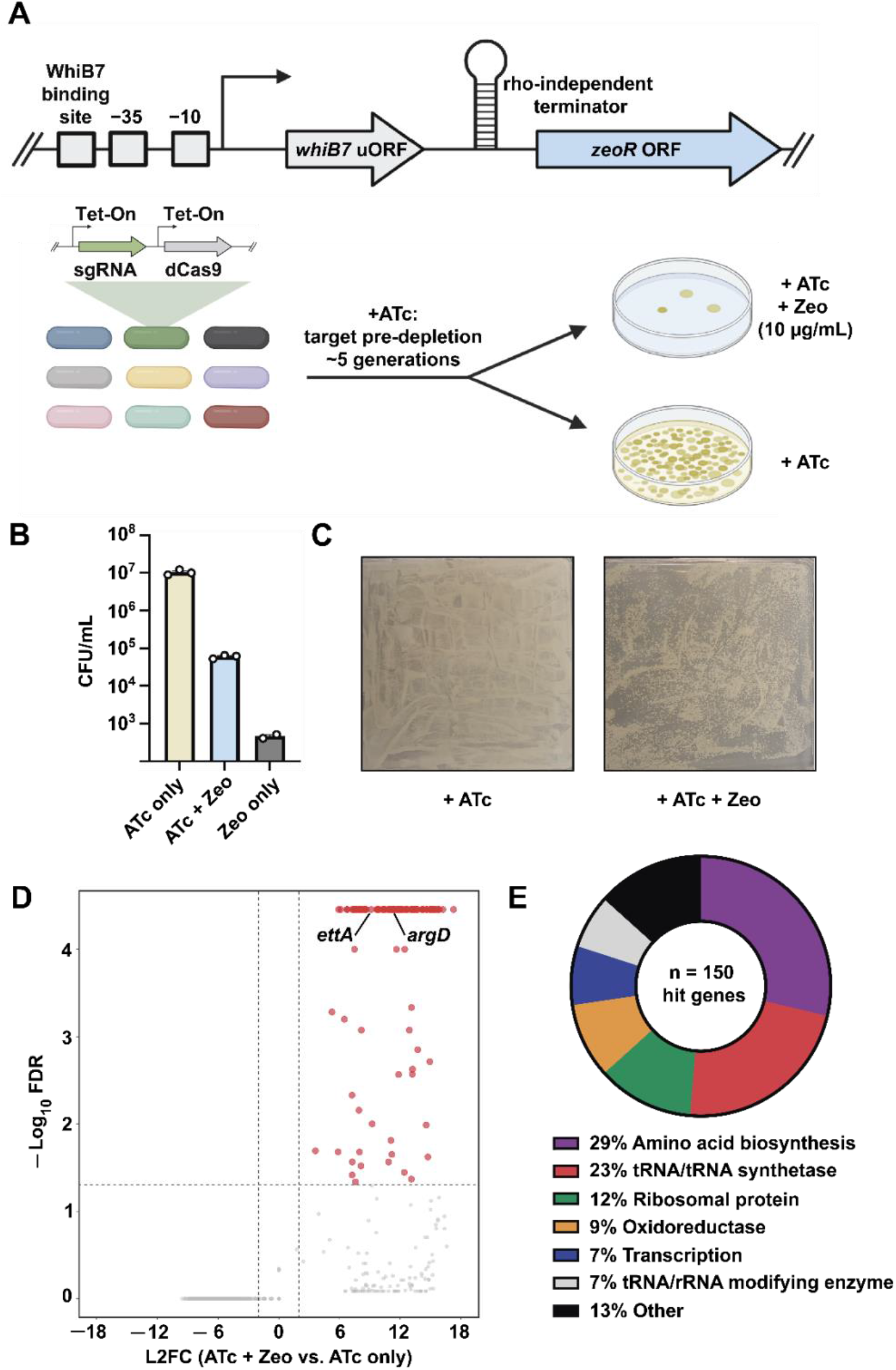
Genome-scale identification of *whiB7*-activating CRISPRi strains in *M. smegmatis*. (A) *whiB7* activation screen workflow. (B,C) CFU quantification (B) and representative plate images (C) from the *M. smegmatis* screen described in panel A. (D) Volcano plot showing log2 fold-change (L2FC) values and false discovery rates (FDR) for each gene in the *whiB7* activation screen. The expected hit genes *ettA* and *argD* are annotated. Hit genes (n=150) were defined as having a L2FC > 2 and FDR < 0.01. (E) Functional categories of hit genes.

This screen identified 150 genes whose inhibition led to enrichment on zeocin containing plates (**Fig. 2D**, **Supplemental Data 1**). Included amongst the 150 hit genes were *ettA* and *argD*, whose inhibition was previously shown to activate *whiB7* expression (**Fig. 1C,E**)^26, 27^. Essential genes were significantly overrepresented amongst hit genes, making up 105 out of 150 hit genes (70%) despite only comprising ∼6% of the *M. smegmatis* genome, highlighting the utility of tunable CRISPRi to investigate phenotypes of essential genes. Amongst the hit genes, the most represented functional categories were genes involved in amino acid biosynthesis, tRNAs, and tRNA synthetases (**Fig. 2E**, **Supplemental Data 1**). Additionally, many hit genes have known or predicted functions in ribosome assembly, maturation, and function. Interestingly, a number of genes involved in transcription initiation, elongation, and termination were identified as hits, possibly reflecting defects in transcription-translation coupling that could perturb uORF translation and formation of the rho-independent terminator, although other mechanisms cannot be excluded^31^. Both succinate dehydrogenase and the cytochrome c oxidase were identified in this screen, potentially consistent with previous work demonstrating the responsiveness of *whiB7* to redox stress^23^. Note that because of the potential polar effect of CRISPRi knockdown, *whiB7* activation could be a result of knockdown specifically of the targeted hit gene, or a downstream gene in a potential operon. Further experiments would be necessary to distinguish direct from polar effects, and thus we present the results of the *whiB7* activation screen without considering potential polar effects.

We next validated several hit genes from the various functional categories in both *M. smegmatis* and *M. tuberculosis* using individual sgRNAs. In accordance with the screen, knockdown of *ettA,* the alanine aminotransferase *aspC,* the transcription antitermination factor *nusB,* and the threonyl-tRNA synthetase *thrS* resulted in strong *whiB7* induction as assessed by qPCR, whereas knockdown of the non-hit essential gene *mmpL3* did not (**Fig. S2A,B**). Using the *PwhiB7:mScarlet* reporter system, we further validated a larger panel of hit and non-hit genes in both *M. smegmatis* (**Fig S2C,D**) and *M. tuberculosis* (**Fig. S2E**). Results with individual strains were largely consistent with the *whiB7* activation screen and concordant between the two bacterial species. Consistent with the interpretation that inhibition of amino acid biosynthetic genes leads to *whiB7* activation as a result of translation stalling within the uORF, we could chemically complement *whiB7* induction in amino acid auxotrophs by provision of the appropriate amino acid (**Fig. S2F**). *whiB7* expression was reversed in an *argD* CRISPRi strain by provision of arginine^26^. Similarly, *whiB7* activation was reversed in the *aspC* knockdown strain by supplying exogenous alanine but not aspartate or glutamate, consistent with prior work demonstrating that contrary to its annotation, *aspC* acts as an alanine aminotransferase^32^.

### Amino acid composition of the uORF dictates the responsiveness of *whiB7* to amino acid starvation

Results from the *M. smegmatis* CRISPRi *whiB7* activation screen suggested that auxotrophy for many different amino acids can result in *whiB7* activation. Biosynthetic genes and/or tRNA/tRNA synthetase pairs related to 18/20 amino acids were identified as hits (**Supplemental Data 1**). Interestingly, genetic (CRISPRi) and pharmacological (6-FABA) inhibition of tryptophan biosynthesis^33^ activated *whiB7* in *M. smegmatis*, but this effect was not seen in *M. tuberculosis* (**Fig. 3A-C)**. This raised the question of what might underly these species- specific differences in *whiB7* activation signals. While the WhiB7 protein sequence is highly conserved across mycobacteria^20^, the *whiB7* uORF displays substantial length and sequence variation across different species (**Fig. 3D, Supplemental Table 1**). Therefore, we interrogated whether coding sequence variation in the uORF could explain the differences in responsiveness to tryptophan deprivation between *M. smegmatis* and *M. tuberculosis*.

**Figure 3:**
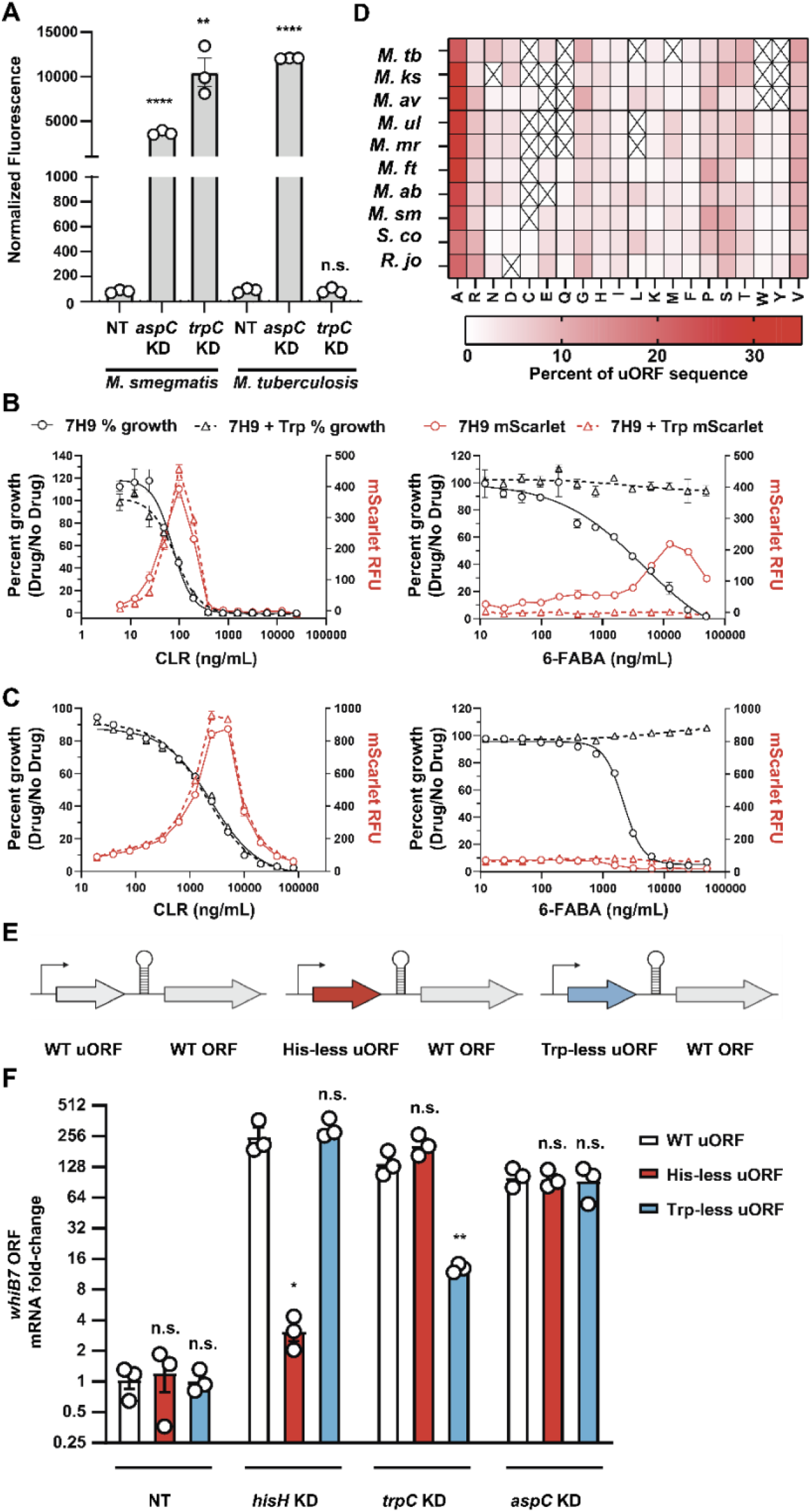
Modulation of the uORF coding sequence alters the response to amino acid deprivation. (A) Normalized fluorescence of the indicated *PwhiB7:mScarlet M. smegmatis* and *M. tuberculosis* CRISPRi strains 24 hours and 8 days after addition of ATc, respectively (mean ± s.e.m., n = 3 replicates). Statistical significance with respect to each non-targeting CRISPRi strain was calculated using a Student’s t-test; **P< 0.01, ****P< 0.0001, n.s. = non-significant. (B-C) Dose response curves (mean ± s.e.m., n = 3 replicates) of the *PwhiB7:mScarlet* reporter strain for clarithromycin (translation inhibitor) and 6-FABA (tryptophan biosynthesis inhibitor), grown in the presence or absence of 1 mM tryptophan. (C) Top row = *M. smegmatis*; (D) bottom row = *M. tuberculosis*. Drug dose-response curves (percent growth) are shown in black and mScarlet fluorescence (RFU) are shown in red. The blue shaded region highlights the differential *whiB7* activation of *M. smegmatis* and *M. tuberculosis* in response to tryptophan limitation by 6-FABA. (D) Amino acid composition (x-axis) of annotated or predicted *whiB7* uORF sequences from the listed bacteria. A black X denotes lack of that amino acid within the indicated *whiB7* uORF. *M. tb* = *M. tuberculosis; M. ks* = *M. kansasii; M. av* = *M. avium; M. ul* = *M. ulcerans; M. mr* = *M. marinum; M. ft* = *M.fortuitum; M. ab* = *M. abscessus; M. sm* = *M. smegmatis; S. co* = *S. coelicolor; R. jo* = *R. jostii*. (E) Genetic architecture of the engineered *M. smegmatis whiB7* uORF variants. (F) *whiB7* ORF mRNA fold-change of the indicated *M. smegmatis* CRISPRi strains 18 hours after addition of ATc (mean ± s.e.m., n = 3 biological replicates). *whiB7* ORF mRNA fold change is relative to *sigA* and normalized to the respective non-targeting CRISPRi strain for each uORF variant. Statistical significance with respect to the WT *whiB7* uORF strain was calculated for each knockdown mutant using a Student’s t-test; *P< 0.05, **P< 0.01, n.s. = non-significant. WT = wild-type.

The *M. smegmatis whiB7* uORF encodes every amino acid except cysteine. In contrast, the *M. tuberculosis* uORF is lacking several amino acids, including tryptophan (**Fig. 3D**). According to the current model of *whiB7* activation^22^, amino acid deficiency should stall uORF translation at specific codons when the ribosome is unable to load the cognate, charged aminoacyl-tRNA. Stalled translation then results in formation of an antiterminator structure and transcriptional readthrough into the *whiB7* ORF^22^. If a particular amino acid is not represented in the uORF (e.g. tryptophan), the inability to produce that amino acid should not stall uORF translation and thus not induce expression of *whiB7*. To test this model, we generated two variants of the *M. smegmatis whiB7* uORF (**Fig. 3E)**. A Trp-less uORF variant was generated where the single tryptophan residue was converted to alanine. A His-less uORF variant had all three histidine residues converted to alanine. None of these mutations are predicted to impact the formation of the *whiB7* antiterminator (**Fig. S3**). We transformed the uORF variants into Δ*whiB7 M. smegmatis* and then monitored the effects of tryptophan, histidine, and alanine deprivation on *whiB7* ORF expression (**Fig. 3F**). Deprivation of tryptophan by knockdown of *trpC* (**Fig. 3F)** or 6-FABA treatment (**Fig. S4A)** induced strong *whiB7* expression in the WT and His-less uORF strains but significantly less *whiB7* expression in the Trp-less uORF strain. Conversely, deprivation of histidine by knockdown of *hisH* induced strong *whiB7* expression in the WT and Trp-less uORF strains but did not induce *whiB7* in the strain harboring the His- less uORF. Importantly, all strains showed high level *whiB7* induction in response to *aspC* knockdown, since all three uORF variants harbored abundant Ala codons. The failure to induce *whiB7* in the uORF variants was not a result of different levels of target gene knockdown (*hisH*, *trpC, aspC*) nor due to differences in basal *whiB7* expression (**Fig. S4A,B**). Taken together, these results are consistent with the model that the ability of *whiB7* to “sense” the depletion of particular amino acids is determined by the amino acid composition of the *whiB7* uORF.

We next analyzed the *whiB7* uORF sequence across ∼50,000 clinical *M. tuberculosis* isolates^27^ to determine whether heterogeneity in this region may influence basal *whiB7* expression or the responsiveness of *whiB7* to amino acid deprivation. Although we did not observe any instances where a mutation introduced a new amino acid not already present in the *whiB7* uORF (**Supplemental Data 2)**, we did identify several rare mutations that led to a robust increase in basal *whiB7* expression (**Fig. S4C**)^34^. These uORF mutations include a loss-of-start codon mutation as well as several distinct indels which likely alter uORF translation dynamics in such a way that promotes transcriptional readthrough into the *whiB7* ORF. We also identified a His22Pro mutation which introduced a di-prolyl motif into the uORF coding sequence that results in constitutive *whiB7* expression. This effect may be explained by ribosome stalling at polyproline motifs^35, 36^, although further experiments are necessary to test this hypothesis. Importantly, none of the *M. tuberculosis* uORF mutations tested in these experiments (**Fig. S3C**) occur within the transcriptional antiterminator or terminator structures, suggesting that the effects of these mutations are due to altered translation dynamics of the uORF and not simply due sequence-based disruption of key RNA secondary structures (**Fig. S3**).

### *whiB7* mediates an adaptive response to alanine starvation

Although *whiB7* expression can be induced by the depletion of many different amino acids, we observed that across actinobacterial species, the *whiB7* uORF is universally rich in alanine (**Fig. 3D**). Depending on the species, 25-40% of the *whiB7* uORF codes for Ala. Each uORF (**Fig. 3D**) was significantly enriched (p < 0.001) for alanine and only alanine relative to the respective proteome of each species^37^. The canonical role of *whiB7* is as an inducible intrinsic resistance mechanism to translation inhibiting antibiotics. What then might account for the strong enrichment for a particular amino acid (Ala) in the *whiB7* uORF? Inspired by classic attenuation mechanisms for *E. coli* tryptophan and leucine biosynthesis^38–40^, as well as the observed responsiveness of *whiB7* to alanine starvation (**Fig. 3F, Fig. S2B-F**), we hypothesized that *whiB7* may play a more general physiological role in regulating alanine metabolism, independent of its canonical role in antibiotic resistance.

To further explore this potential physiological role for *whiB7*, we performed a genome-wide differential vulnerability screen (**Fig. 4A**)^30^. Differential vulnerability screens identify genes that become more or less sensitive to CRISPRi inhibition between two (or more) genetic backgrounds or growth conditions^30^. To perform this screen, the same tunable sgRNA library^30^ used to identify upstream *whiB7* activators (**Fig. 2A**) was transformed into Δ*whiB7 M. smegmatis*^26^. The resulting CRISPRi library was then passaged for approximately 30 generations in the presence or absence of ATc. Every 2.5 or 5 generations, we harvested genomic DNA and analyzed sgRNA abundance by deep sequencing. We then compared gene vulnerability between wild-type *M. smegmatis*^30^ and the Δ*whiB7* mutant. Quantification of gene vulnerability revealed strong concordance between the two strains (R^2^ = 0.963), although there were rare differentially vulnerable genes. We identified a total of 20 differentially vulnerable genes, 9 of which were more vulnerable in Δ*whiB7* (ΔVI < –2) and 11 of which were less vulnerable (ΔVI > 2) in Δ*whiB7*.

**Figure 4:**
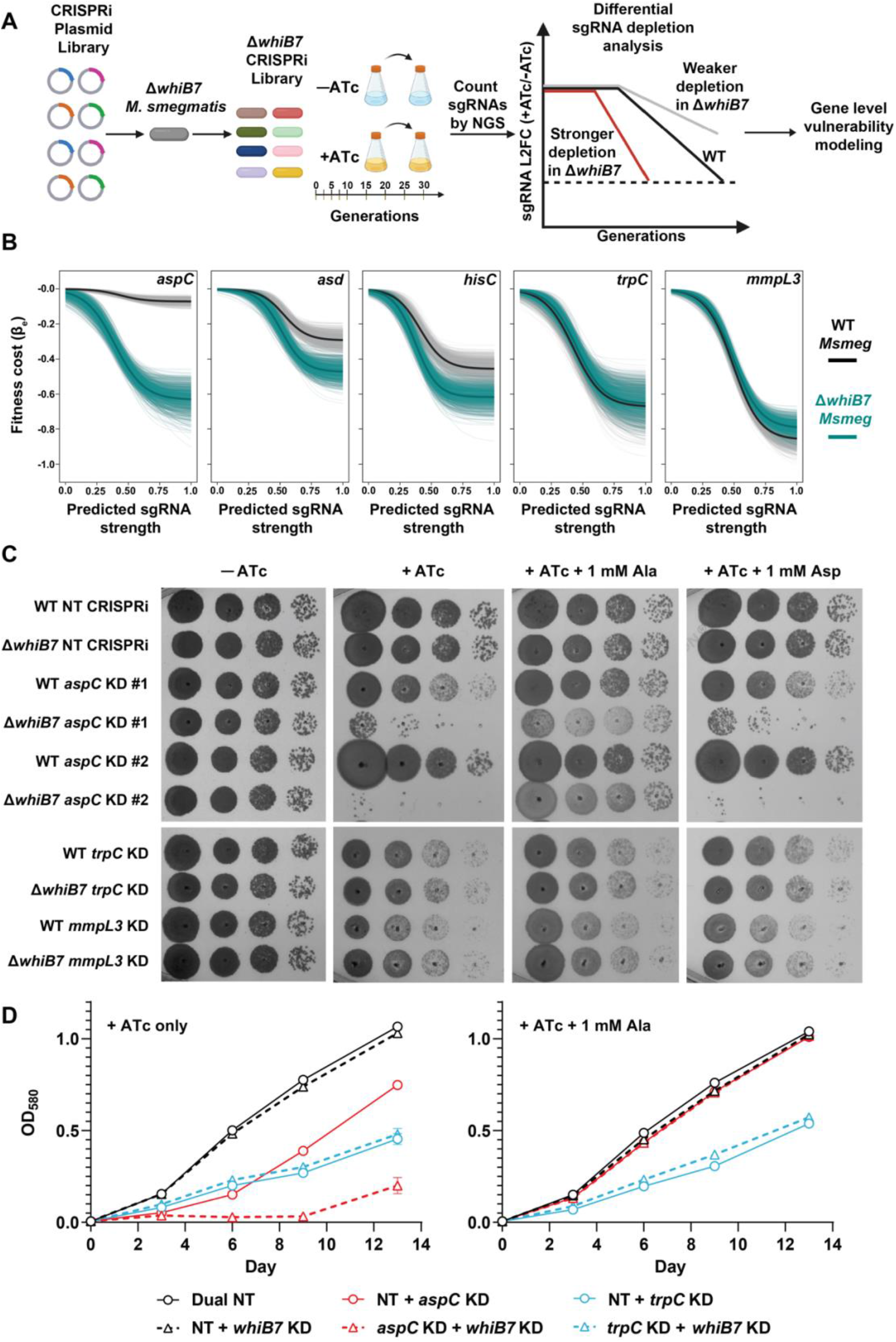
Identification of differential vulnerabilities in Δ*whiB7*. (A) Differential vulnerability screen in Δ*whiB7 M. smegmatis.* This screen identifies genes that become more or less sensitive to CRISPRi inhibition between wild-type and Δ*whiB7 M. smegmatis*. Please see the Materials and Methods section and Bosch et al.^30^ for further details on screen analysis. (B) Expression-fitness relationships for the five indicated *M. smegmatis* genes. The fitness cost (beta_E) is plotted as a function of predicted sgRNA strength (an estimate of the magnitude of target knockdown). Both *aspC, asd,* and *hisC* are more vulnerable in Δ*whiB7 M. smegmatis* (tourquise). (C) Growth of the indicated *M. smegmatis* CRISPRi strains monitored by spotting serial dilutions of each strain on the indicated media. Supplemental alanine and aspartate were added at 1 mM. Note that hypomorphic sgRNAs that are predicted to lead to intermediate levels of knockdown are shown for *trpC* and *mmpL3*, as strong sgRNAs leading to high-level knockdown would block growth of both wild-type and Δ*whiB7 M. smegmatis* and not be relevant controls for differential vulnerability. (D) Growth of the indicated *M. tuberculosis* dual-gene knockdown CRISPRi strains in 7H9 + ATc, with or without supplemental alanine (1 mM). Dual NT represents a CRISPRi plasmid encoding two non- targeting sgRNAs; NT + *whiB7* KD represents a CRISPRi plasmid encoding a single non-targeting sgRNA and a *whiB7* targeting sgRNA; *aspC* KD + *whiB7* KD represents a CRISPRi plasmid encoding one sgRNA targeting *aspC* and a separate sgRNA targeting *whiB7*.

The gene which showed the greatest difference in vulnerability between WT and Δ*whiB7* was the alanine aminotransferase *aspC*, although more subtle differences were also observed for aspartate semialdehyde dehydrogenase *asd* and histidinol-phosphate aminotransferase *hisC* (**Fig. 4B**). All three of these genes were also identified as genes whose inhibition resulted in *whiB7* activation (**Fig. 3A,F**, **Fig. S2A-E, Supplemental Data 1**). The differential vulnerability screen indicated that silencing of *hisC, asd,* and particularly *aspC* resulted in a much stronger fitness costs in Δ*whiB7* than wild-type *M. smegmatis*, potentially consistent with our hypothesis that *whiB7* plays a physiological role in amino acid metabolism. Interestingly, the vast majority of other amino acid biosynthetic genes and essential processes were not differentially vulnerable (**Supplemental Data 3**, **Fig. 4B**), suggesting that *whiB7* is specifically required to maintain fitness under physiological conditions in which AspC and to a lesser extent HisC and Asd activity is perturbed.

Using individual CRISPRi strains, we confirmed that *aspC* and *asd* were substantially more vulnerable in Δ*whiB7* compared to a wild-type *M. smegmatis* strain (**Fig. 4C, Fig. S5B**). These phenotypes were not a result of a CRISPRi polar effect, as the potential operonic genes downstream of both *aspC* and *asd* were not differentially vulnerable. This effect was specific, as silencing of the non-hit control genes *trpC* and *mmpL3* was not associated with a greater fitness cost in Δ*whiB7* (**Fig. 4C**). The fitness defect could be reversed in the *aspC* CRISPRi strains by supplementing exogenous alanine, but not aspartate (**Fig. 4C, Fig. S2F**)^32^. Interestingly, amino acid supplementation exacerbated the growth defect associated with *asd* knockdown in both the wild-type and Δ*whiB7* (**Fig. S5C**), which may reflect the accumulation of toxic intermediates of aspartate metabolism that cannot be eliminated in the absence of Asd. We next validated *aspC* differential vulnerability in *M. tuberculosis*. Concurrent knockdown of both *aspC* and *whiB7* resulted in stronger growth inhibition than knockdown of *aspC* alone. This effect could be reversed by alanine supplementation, but not glutamate or aspartate (**Fig. 4D, Fig. S5D**). Knockdown of *whiB7* did not exacerbate the growth defect associated with *trpC* knockdown, demonstrating the specificity of this effect for *aspC*.

Why is *aspC* more vulnerable in Δ*whiB7*? Previous RNA sequencing and ChIP sequencing data suggests that *aspC* is part of the *whiB7* regulon in *M. smegmatis*^17^*, M. abscessus*^17, 41^, and *S. coelicolor*^42^. Consistent with these results, we were able to confirm the interaction of WhiB7 with the *M. smegmatis aspC* promoter by chromatin immunoprecipitation (ChIP) RT-qPCR (**Fig. 5A**). These observations, combined with the increased vulnerability of *aspC* in Δ*whiB7*, suggest that *whiB7* and *aspC* may participate in a feedback loop. In this model, alanine depletion results in translational stalling in the alanine-rich *whiB7* uORF. This results in the induction of WhiB7 and WhiB7-mediated upregulation of *aspC*, thus restoring alanine levels. In the absence of *whiB7*, alanine depletion is not sensed, *aspC* is not induced, and the cell fails to initiate an adaptive response to restore alanine levels. To test this model, we first targeted *aspC* by CRISPRi and measured the level of *aspC* knockdown in both WT and Δ*whiB7* strains. If WhiB7 is important to upregulate *aspC* expression under conditions of alanine starvation, then *aspC* knockdown should be stronger in the Δ*whiB7* strain. Consistent with the model, we observed significantly greater *aspC* knockdown in the Δ*whiB7* knockout strain (**Fig. 5B**). No differences in target knockdown were observed for the control genes *mmpL3* or *trpC,* indicating that CRISPRi efficiency does not differ between the wild-type and Δ*whiB7*. Also consistent with the model, we observed that *ettA* knockdown strains, which serve as a model for robust, constitutive *whiB7* activation^27^, show increased expression of *aspC* (**Fig. 5C**). Induction of *aspC* in the *ettA* knockdown strains could specifically be reversed by knocking down or knocking out *whiB7*, in accordance with prediction that *whiB7* is critical for *aspC* induction (**Fig. 5C, D**). Taken together, these data support the conclusion that *whiB7* plays an important role outside of canonical intrinsic drug resistance by regulating cellular alanine levels through a feedback loop with *aspC* (**Fig. 5E**).

**Figure 5:**
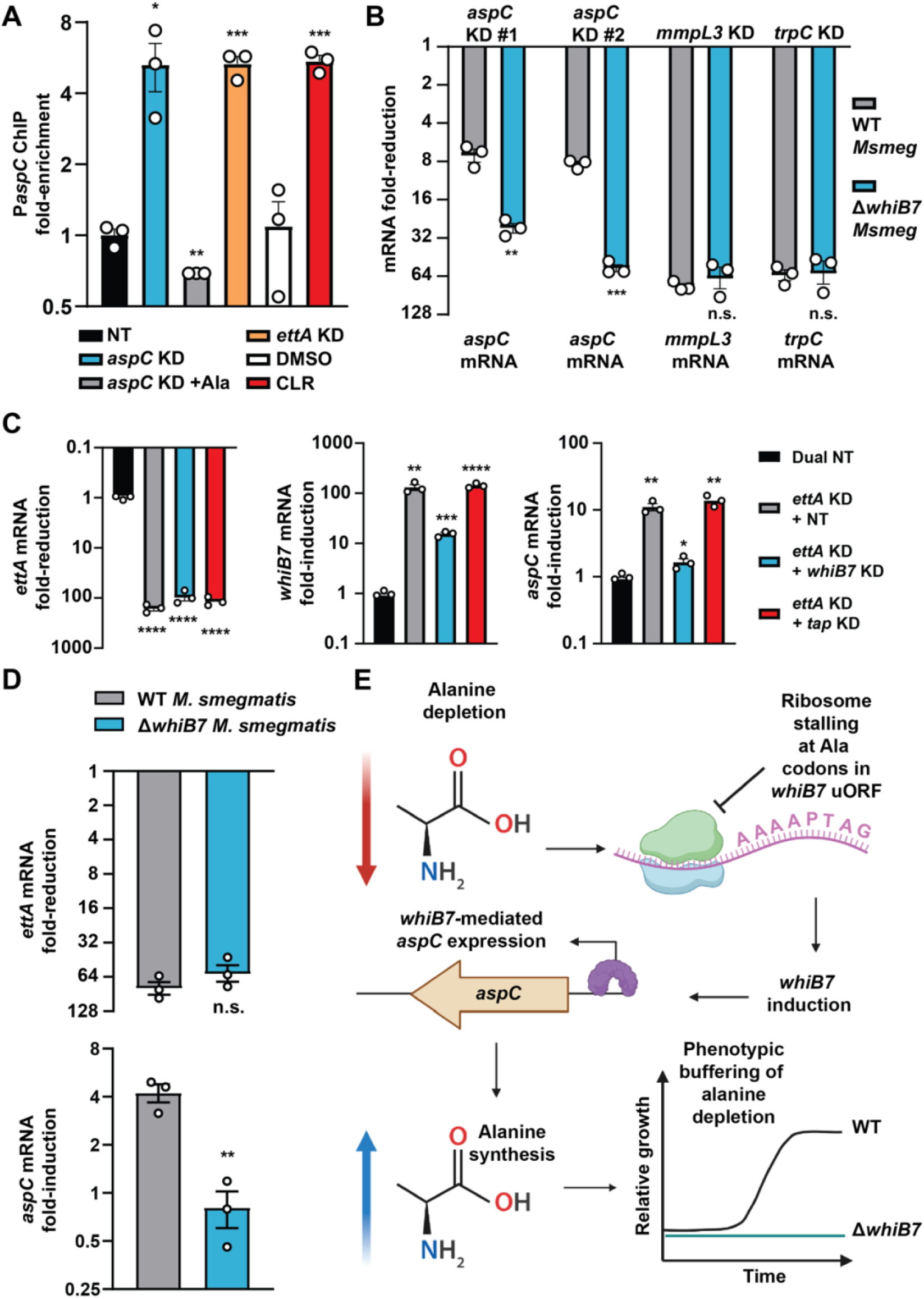
*whiB7* coordinates a feedback loop with *aspC*. (A) ChIP RT-qPCR of the *aspC* promoter in the indicated *M. smegmatis* WhiB7 N-terminal 3X-FLAG strains. Fold-enrichment of *aspC* promoter qPCR signal relative to the control *trpC* promoter (WhiB7- independent) is indicated. *whiB7* was activated either by CRISPRi knockdown of *ettA* or *aspC* (18 hours +ATc), or treatment with clarithromycin for 12 hours. Statistical significance with respect to the non-targeting CRISPRi strain or DMSO control was calculated using a Student’s t-test, **P< 0.01, ***P<0.001 n.s. = non-significant. (B) Relative mRNA levels of the indicated genes in the indicated CRISPRi strains 15 hours after addition of ATc (mean ± s.e.m., n = 3 biological replicates). mRNA fold-change for the indicated gene at the bottom of each pair of bar graphs was calculated relative to *sigA* and normalized to the respective non-targeting CRISPRi strain for WT *M. smegmatis* or the Δ*whiB7* strain. Grey = WT, blue = Δ*whiB7.* Statistical significance between the WT and Δ*whiB7* strain was calculated using a Student’s t-test, **P< 0.01, ***P<0.001. (C) Relative mRNA levels of the indicated genes in the indicated CRISPRi strains 5 days after addition of ATc (mean ± s.e.m., n = 3 biological replicates) in *M. tuberculosis*. mRNA fold-change for the gene indicated on the y-axis was calculated relative to *sigA* and normalized to the respective non-targeting CRISPRi dual non-targeting CRISPRi strain. Statistical significance for each strain was calculated with respect to the dual non-targeting (NT) strain for each of the indicated genes using a Student’s t-test, *P<0.05, **P< 0.01, ***P<0.001, ****P<0.0001. (D) Relative mRNA levels of the indicated genes in *ettA* knockdown CRISPRi strains 18 hours after addition of ATc (mean ± s.e.m., n = 3 biological replicates). mRNA fold-change for the gene indicated on the y-axis was calculated relative to *sigA* and normalized to the respective non-targeting CRISPRi strain for WT *M. smegmatis* or the Δ*whiB7* strain. Grey = WT, blue = Δ*whiB7.* Statistical significance between the WT and Δ*whiB7* strain was calculated using a Student’s t-test, **P< 0.01. (E) Proposed model for the *whiB7*-*aspC* feedback loop.

## DISCUSSION

*whiB7*-mediated drug resistance complicates treatment of mycobacterial diseases^15, 43–45^. Here, we took an functional genomics approach to globally define the upstream signals that trigger *whiB7* activation, as well as downstream physiological processes that depend on *whiB7*. We discovered 150 genes whose inhibition leads to constitutive *whiB7* activation. Hit genes include expected processes critical for translation, including amino acid biosynthesis, tRNAs, tRNA synthetases, ribosome assembly, maturation, and function. We also identified numerous unexpected hit genes involved in diverse processes including transcription, central carbon metabolism, the electron transport chain, and uncharacterized genes. It will be important to determine how silencing of these unexpected hit genes results in *whiB7* activation. Essential genes were significantly overrepresented amongst hit genes, highlighting the ability of tunable CRISPRi to investigate phenotypes of essential genes and more broadly demonstrating the utility of CRISPRi reporter screens to identify genes that functionally interact with a regulatory sequence. Note that not all essential genes whose inhibition activates *whiB7* may be recovered in the *PwhiB7:zeoR* screen. Essential hit genes require sgRNAs strong enough to reduce gene product activity sufficiently to induce *whiB7*, but not reduce fitness enough to prevent colony formation. Such false negatives, likely enriched for small genes for which fewer sgRNAs are available, may be recoverable using the *PwhiB7:mScarlet* reporter coupled with FACs to obviate the need for colony outgrowth. Downstream of *whiB7*, we identified a critical feedback loop between *whiB7* and *aspC* that coordinates an adaptive response to alanine deprivation. AspC has no known role in promoting intrinsic drug resistance^27^, thus demonstrating that *whiB7* plays an important physiologic role outside of its canonical role in intrinsic drug resistance by regulating cellular alanine levels.

Our *whiB7* activation screen results are consistent with the model that *whiB7* senses general translation stress via the uORF^22^. Our results build on this model of uORF-mediated translation sensing by showing that the specific amino acid composition of the uORF determines the sensitivity of *whiB7* activation to amino acid deprivation. This more nuanced model suggests that species-specific differences in uORF composition may produce unique *whiB7* activation patterns during stress conditions. This may be particularly informative for drug discovery efforts targeting amino acid biosynthetic enzymes or aminoacyl tRNA synthetases^33, 46, 47^. Drugs which activate *whiB7* will likely antagonize aminoglycosides, macrolides, and potentially other classes of antibiotics^15^. Thus, in order to avoid antagonistic drug combinations, new antimycobacterial compounds would ideally avoid activation of the *whiB7* pathway^15^. Accordingly, targeting leucine or tryptophan metabolism in *M. tuberculosis* will likely result in minimal *whiB7* induction, whereas targeting histidine or alanine metabolism will likely result in high level *whiB7* induction. The results presented here will be informative for predicting and avoiding these types of *whiB7*-mediated antagonistic drug interactions.

Although the *whiB7* uORF displayed substantial sequence diversity across actinobacterial species, the conserved enrichment for alanine suggested that *whiB7* might play a physiological role in alanine metabolism. Indeed, we found that the alanine biosynthetic enzyme *aspC* becomes substantially more vulnerable in the absence of *whiB7*, and we present evidence that *whiB7* is capable of sensing alanine levels and buffering *aspC* knockdown via a feedback loop. Our data suggest a similar mechanism may operate for *asd* and the *hisC* operon, although future work is necessary to test this hypothesis. The model presented here shows parallels to the classic attenuation mechanisms described in *E. coli* amino acid biosynthetic operons. Several *E. coli* operons such as the tryptophan and leucine operons have a leader/uORF sequence rich in the corresponding amino acid ^38–40^. Deprivation of that particular amino acid results in ribosome stalling in the leader peptide, resulting in antitermination and transcriptional readthrough into the downstream biosynthetic genes to restore amino acid levels. While the mycobacterial *whiB7* uORF typically encodes a longer and more diverse peptide sequence that can broadly sense amino acid starvation, it appears to be particularly tuned to sense alanine deprivation. Why *whiB7* evolved such an “Ala leader” mechanism tuned to sense and specifically regulate alanine levels, as opposed to some other amino acid, remains an interesting and open question. Given the centrality of alanine in bacterial metabolism, we cannot rule out functions for this feedback loop beyond the proteinogenic functions for alanine. For example, flux through AspC could enable alanine to serve as an indirect readout of pyruvate levels and the various metabolic pathways in which pyruvate participates. Further, L-alanine is a direct precursor to D-alanine, a core component of cell wall peptidoglycan. Future studies may investigate the broader metabolic and physiological consequences of *whiB7*-mediated *aspC* regulation.

Lastly, the results presented here may help further our understanding of the complex interactions between mycobacterial metabolism, the host immune response and antibiotic activity. In the context of TB, immune activation has been shown to induce drug tolerance^48, 49^. Although the molecular basis for this phenomenon is likely to involve many distinct host and bacterial pathways, *whiB7* induction may be an important contributor to host-induced drug tolerance^19, 26^. Previous work showed that *M. tuberculosis whiB7* is upregulated upon infection of both resting and IFN-γ-activated macrophages^28, 50^, which may promote rifampicin and aminoglycoside tolerance mediated by the Tap efflux pump^19^. Our results may help define the host-imposed stressors and affected bacterial pathways that lead to intramacrophage *whiB7* induction. Future studies could use the *M. tuberculosis PwhiB7:mScarlet* reporter to dissect *whiB7* expression patterns across a diverse set of host immune cells including different subsets of macrophages, neutrophils, and dendritic cells, as well as scenarios involving drug treatment^51, 52^. We hope that these studies will shed light on how distinct intracellular environments influence mycobacterial metabolism, thus shaping drug resistance and tolerance patterns.

## Supporting information

Supplemental_Data_1_whiB7_zeo_screen_results

Supplemental_Data_2_whiB7_uORF_variants

Supplemental_Data_3_whiB7KO_vulnerability

Supplemental_file_plasmids_primers

## ACKNOWLEDGMENTS

We thank members of the Rock laboratory and Alexandre Gouzy for comments on the manuscript and/or helpful discussions, Sarah Schrader, Julien Vaubourgeix, and Carl Nathan for sharing the Δ*whiB7 M. smegmatis* strain, Chendong Pan, Xing Wang, Jenny Xiang, and Adrian Tan of the Weill Cornell Genomics Core for help with NGS, Connie Zhao of the Rockefeller University Genomics Resource Center for help with chromatin immunoprecipitation, and Sang Hyun Cho and Scott Gary Franzblau for sharing ecumicin. We thank Elizabeth Campbell, Mira Lilic, and Maddie Delbeau for experimental guidance and for helpful discussions. This work was supported by the Potts Memorial Foundation (M.A.D.), a joint NIH tuberculosis research units network (TBRU - N) grant (U19AI162584, J.M.R), and an NIH/NIAID New Innovator Award (1DP2AI144850-01, JMR). The illustrations in Figures 1A, 2A, 4A, and 5E were generated using BioRender software.

## AUTHOR CONTRIBUTIONS

Conceptualization: N.C.P. and J.M.R. Investigation: N.C.P., C.R., V.M-G., S.L., and K.M.L. Data Analysis: N.C.P., M.A.D, and Z.A.A., Writing—original draft: N.C.P. and J.M.R. Writing—review and editing: N.C.P., S.L, Z.A.A., M.A.D., and J.M.R. Funding acquisition: J.M.R. Supervision: J.M.R.

## MATERIALS AND METHODS

### Mycobacterial cultures

*M. smegmatis* strains are derivatives of the mc^2^ 155 strain. *M. abscessus* strains are derivatives of the ATCC19977 strain. *M. tuberculosis* strains are derivatives of the H37Rv background. All mycobacteria were grown at 37°C in Difco Middlebrook 7H9 broth or on 7H10 agar supplemented with 0.2% glycerol (7H9) or 0.5% glycerol (7H10), 0.05% Tween-80 and either 1X albumin-dextrose-catalase (ADC) for *M. smegmatis* or 1X oleic acid-albumin-dextrose-catalase (OADC) for *M. tuberculosis* and *M. abscessus*. For CRISPRi experiments, anhydrotetracycline (ATc) was used at 100 ng/ml. For plasmid selection kanamycin was used at 20 µg/mL and zeocin was used at 20 µg/mL. For the *whiB7* induction screen in *M. smegmatis* zeocin was used at 10 µg/mL. *M. tuberculosis* cultures were grown standing in tissue culture flasks (unless otherwise indicated) with 5% CO_2_, whereas *M. smegmatis* strains were grown in shaking conditions in sealed culture vessels. Amino acids were supplemented to the media at the indicated concentration.

Relative growth of individual CRISPRi strains was determined by spotting assay. Ten-fold serial dilutions (starting at 30,000 cells/spot) were plated on 7H10 with or without 100 ng/mL ATc. Plates were incubated at 37°C and imaged after 2 days.

### P*whiB7:mScarlet* reporter assays

mScarlet fluorescence was measured using a Tecan spark plate reader with an excitation of 563 nm and and emission of 600 nm. For *M. smegmatis* assays, plates were cultured in the plate reader for 48 hours with shaking at 500 rpm. Fluorescence and optical density were measured every 30 minutes. For *M. tuberculosis* assays, plates were incubated under standing conditions and were read for fluorescence and optical density at 4 to 7 days post-plating. Normalized fluorescence was calculated by dividing the background-adjusted fluorescence value by the background-adjusted optical density value.

### Antibacterial activity measurements

All compounds (see supplemental table 2) were dissolved in DMSO (VWR V0231) and dispensed using an HP D300e digital dispenser in a 384-well plate format using a 2-fold dilution series. DMSO did not exceed 1% of the final culture volume and was maintained at the same concentration across all samples. Cultures were growth synchronized to late logarithmic phase (∼ OD_600_ = 0.8) and then back-diluted to a starting OD_600_ of 0.01. 50 µl cell suspension was plated in triplicate in wells containing the test compound. Plates were incubated standing at 37 °C with 5% CO2. OD_580_ was evaluated using a Tecan Spark plate reader at 10–11 days post-plating and percent growth was calculated relative to the DMSO vehicle control for each strain. IC_50_ measurements were calculated using a nonlinear fit in GraphPad Prism. For all MIC curves, data represent the mean ± s.e.m. for triplicates. Data are representative of two independent experiments.

### *whiB7* activation CRISPRi screen

The *M. smegmatis* strain harboring the P*whiB7:zeoR* construct was transformed with the genome-wide CRISPRi library described in Bosch et al.^30^ (Addgene 163955) with greater than 400-fold coverage. Transformants were plated on complete 7H10 agar with kanamycin at 20 µg/mL. After 3 days of growth, biomass was collected in 10% glycerol and single cell suspensions were created using the GentleMACS dissociator (M tube, 2X RNA cycle). After passaging in complete 7H9 and filtering through a pluriselect filter (10 µm) stocks glycerol stocks were frozen. For the screen one stock of the CRISPRi library was thawed into 25 mL of complete 7H9. After growth to late log-phase, 3 parallel cultures were started at an OD_600_ of 0.025 in the presence of ATc. After 16 hours of pre-depletion, each culture was diluted to 1x10^7^ CFU was plated onto plates containing ATc with or without 10 µg/mL zeocin. In parallel, a non-ATc treated library culture was plated at the same density on 10 µg/mL zeocin without ATc. In parallel, serial dilutions of pre-depleted and non pre-depleted cultures were plated on smaller agar plates for CFU enumeration. All plates were counted and harvested after 3 days outgrowth. Biomass was from each plate was scraped into TE buffer and subjected to GentleMACS dissociation (M tube, 2X RNA cycle). Approximately 50 OD_600_ units were then subjected to gDNA extraction via the TE-lysozyme method described below.

### Δ*whiB7* vulnerability CRISPRi screen

The Δ*whiB7 M. smegmatis* strain described in Schrader et al.^26^, was transformed with the genome-wide CRISPRi library (Addgene 163955) with greater than 200-fold coverage. Transformants were plated on complete 7H10 agar with kanamycin at 12.5 µg/mL, due to potential increase in kanamycin sensitivity in the absence of *whiB7*. After 3 days of growth, biomass was collected and single cell suspensions were generated using the same process as the activation screen (see above). For the passaging time course, one stock of the CRISPRi library was thawed and expanded into 25 mL of complete 7H9 + kanamycin 12.5 µg/mL. Parallel, triplicate cultures were set up with or without ATc at 100 ng/mL. At each indicated timepoint (generations), cultures were harvested for genomic DNA isolation. Remaining culture was used for continued passaging. Each sample was then subjected to gDNA extraction via the TE-lysozyme method described below.

### Genomic DNA extraction

Genomic DNA was isolated from bacterial pellets using the CTAB-lysozyme method as previously described^30^. After drug treatment 10-20 OD_600_ units of the cultures were pelleted by centrifugation (10 minutes at 4,000xg) and were resuspended in 1ml PBS + 0.05% Tween-80. Cell suspensions were centrifuged again for 5min at 4,000×g, the supernatant was removed, and pellets were frozen until processing. For the DMSO-treated culture and the cultures treated with supra-MIC drug concentrations, 500 µL of the remaining culture was spread evenly on a 15 cm petri dish containing complete 7H10 + 0.4% activated charcoal. After 17 -21 days of outgrowth, the biomass was scraped off of the plate using PBS + 0.05% Tween-80 and gDNA was processed the identically to the pellets obtained directly from liquid culture. Upon thawing, pellets were resuspended in 800 µl TE buffer (10mM Tris pH 8.0, 1mM EDTA) + 15mg/mL lysozyme (Alfa Aesar J60701-06) and incubated at 37 °C for 16h. Next, 70µl 10% SDS (Promega V6551) and 5 µl proteinase K (20 mg/mL, Thermo Fisher 25530049) were added and samples were incubated at 65°C for 30min. Subsequently, 100 µl 5M NaCl and 80 µl 10% CTAB (Sigma Aldrich H5882) were added, and samples were incubated for an additional 30min at 65 °C. Finally, 750 µl ice - cold chloroform was added and samples were mixed. After centrifugation at 16,100×g and extraction of the aqueous phase, samples were removed from the biosafety level 3 facility. Samples were then treated with 25 µg RNase A (Bio Basic RB0474) for 30min at 37 °C, followed by extraction with phenol:chloroform:isoamyl alcohol (pH 8.0, 25:24:1, Thermo Fisher BP1752I-400), then chloroform. Genomic DNA was precipitated from the final aqueous layer (600 µl) with the addition of 10 µl 3M sodium acetate and 360 µl isopropanol. DNA pellets were spun at 21,300×g for 30min at 4 °C and washed 2× with 750 µl 80% ethanol. Pellets were dried and resuspended with elution buffer (Qiagen 19086) before spectrophotometric quantification. The concentration of isolated genomic DNA was quantified using the DeNovix dsDNA high sensitivity assay (KIT-DSDNA-HIGH-2; DS-11 series spectrophotometer/fluorometer).

### Library preparation for Illumina sequencing of CRISPRi libraries

Next generation sequencing was performed as follows. The unique barcoded region was amplified from 500ng genomic DNA with 16 cycles of PCR using NEBNext Ultra II Q5 master mix (NEB M0544L) as described in^30^. Each PCR reaction contained a unique indexed forward primer (0.5μM final concentration) and a unique indexed reverse primer (0.5μM). Forward primers contain a P5 flow cell attachment sequence, a standard Read1 Illumina sequencing primer binding site and custom stagger sequences to guarantee base diversity during Illumina sequencing. Reverse primers contain a P7 flow cell attachment sequence, a standard Read2 Illumina sequencing primer binding site and unique barcodes to allow for sample pooling during deep sequencing. Following PCR amplification, each ∼230bp amplicon was purified using AMPure XP beads (Beckman –Coulter A63882) using one-sided selection (1.2×). Bead-purified amplicons were further purified on a Pippin HT 2% agarose gel cassette (target range 180–250bp; Sage Science HTC2010) to remove carry-over primer and genomic DNA. Eluted amplicons were quantified with a Qubit 2.0 fluorometer (Invitrogen), and amplicon size and purity were quality controlled by visualization on an Agilent 2100 bioanalyzer (high sensitivity chip; Agilent Technologies 5067–4626). Next, individual PCR amplicons were multiplexed into 10nM pools and sequenced on an Illumina sequencer according to the manufacturer’s instructions. To increase sequencing diversity, a PhiX spike-in of 2.5–5% was added to the pools (PhiX sequencing control v3; Illumina FC-110- 3001). Samples were run on the Illumina NovaSeq 6000 platform (single-read 1 ×85 cycles and 6 × i7 index cycles).

### Analysis of *whiB7* activation screen

Sequencing counts were obtained in the manner described by^30^. Counts were normalized for sequencing depth and an sgRNA limit of detection (LOD) cut-off was set at 20 counts in the ATc only condition. Only sgRNAs that made the LOD cut-off (i.e. counts > 20) were analyzed further. sgRNA counts were analyzed using MAGeCK ^53^ (version 0.5.9.2) in python (version 2.7.16) comparing each + zeocin + ATc condition to the matched control (+ATc only) sample. Gene-level log2 fold change (L2FC) was calculated using the ‘alphamedian’ approach specified with the ‘gene-lfc-method’ parameter, which estimates the gene-level L2FC as the median of sgRNAs that are ranked above the default cut off in the Robust Rank Aggregation used by MAGeCK. Negative control sgRNAs were used to calculate the null distribution and to normalize counts using the ‘-- control-sgrna’ and ‘– normalization control’ parameters, respectively. MAGeCK gene summary output results can be found in Source Data 1. A gene was determined to be an enriched hit if it had a false discovery rate (FDR) < 0.01 and a log2 fold change (L2FC) > 2 in positive selection.

### Differential vulnerability analysis of Δ*whiB7* CRISPRi screen

Gene vulnerability in the WT and ΔwhiB7 *M. smegmatis* strains was determined using the vulnerability model previously described^30^ with some modifications. Briefly, under the updated model, read counts for a given sgRNA in the minus ATc conditions were modeled using a Negative Binomial distribution with a mean proportional to the counts in the plus ATc condition, plus a factor representing the log2 fold-change:

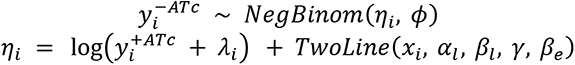

where *λ*_*i*_ is an sgRNA-level correction factor estimated by the model, *x*_*i*_ represents the generations analyzed for the i-th guide, and the TwoLine function represents the piecewise linear model previously described, which models sgRNA behavior over time^30^. The logistic function describing gene-level vulnerabilities was simplified by setting the starting point of the curve (K) equal to 0, representing the fact that weakest possible sgRNAs are expected to impose approximately no effect on bacterial fitness i.e.:

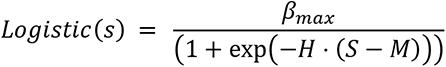

The Bayesian vulnerability models were run for each condition independently, and samples for all the parameters were obtained using Stan (4 chains with 1,000 warmup iterations, and 4000 samples total). Differential vulnerabilities were estimated by taking the difference of pairwise (guide-level) vulnerability estimates, resulting in posterior samples of the differential vulnerability expected for a gene, between the two strains.

### RNA extraction and RT-qPCR

Total RNA extraction was performed as previously described^30^. Briefly, ∼2 OD_600_ units of bacteria were added to an equivalent volume of GTC buffer (5 M guanidinium thiocyanate, 0.5% sodium N-lauroylsarcosine, 25 mM trisodium citrate dihydrate and 0.1 M 2-mercaptoethanol), pelleted by centrifugation, resuspended in 1 ml TRIzol (Thermo Fisher 15596026) and lysed by zirconium bead beating (MP Biomedicals 116911050). Chloroform (0.2 ml) was added to each sample and phases were separated by centrifugation. The aqueous phase was then purified by Direct-zol RNA miniprep (Zymo Research R2052). Residual genomic DNA was removed by TURBO DNase treatment (Invitrogen Ambion AM2238). After RNA cleanup and concentration (Zymo Research R1017), 3 µg RNA per sample was reverse transcribed into complementary DNA (cDNA) with random hexamers (Thermo Fisher 18-091-050) following the manufacturer’s instructions. RNA was removed by alkaline hydrolysis and cDNA was purified with PCR cleanup columns (Qiagen 28115). Next, knockdown or induction of target genes was quantified by SYBR green dye-based quantitative real-time PCR (Applied Biosystems 4309155) on a Quantstudio System 5 (Thermo Fisher A28140) using gene-specific qPCR primers (5 µM), normalized to sigA (rv2703) and quantified by the ΔΔCt algorithm. All gene-specific qPCR primers were designed using the PrimerQuest tool from IDT (https://www.idtdna.com/PrimerQuest/Home/Index) and then validated for efficiency and linear range of amplification using standard qPCR approaches. Specificity was confirmed for each validated qPCR primer pair through melting curve analysis. All qPCR primers used in this study can be found in the Supplemental Plasmids and Primers Table.

### Chromatin immunoprecipitation (ChIP) RT-qPCR

Chromatin immunoprecipitation of an N-terminally 3X-FLAG-tagged WhiB7 was performed according to the protocol described by Jaini et al^54^. Briefly, merodiploid strains expressing a second copy of WhiB7 (driven by the native promoter), either tagged or untagged, as well as the CRISPRi sgRNA of interest were cultured + ATc for 18 hours and grown to late log phase (OD_600_ ∼0.7) to allow for induction of *whiB7*. Alternatively, cultures (no CRISPRi) were treated with either DMSO or clarithromycin (to induce *whiB7*) for 12 hours. Protein-DNA cross- linking was carried out by adding formaldehyde to the cultures to a final volume of 1% and shaking at room temperature for 30 minutes. Cross-linking was quenched by the addition of glycine (250 mM final). Cells were pelleted and mechanically lysed by bead-beating (max speed) 3-times for 30 seconds (1 minute pause between cycles) in the presence of protease inhibitor. Clarified cell lysates (500 µL) were subjected to sonication using the Covaris S220 (Power = 140, Duty = 5, Cycles/Burst = 200) for 18 minutes with microTUBE 500 AFA tubes (Covaris 520185) and each sample was incubated with 5 µg anti-FLAG M2 antibody (Millipore A2220) at 4°C overnight. Immune complexes were purified using 100 µL of protein G agarose slurry (Thermo Scientific 20398) for 2 hours. After washing, DNA protein complexes were eluted at 65°C with Tris-EDTA buffer + 1% SDS, 2 times and the separate eluates were pooled. Reverse cross-linking was performed with 1 mg/mL (final) proteinase K at 37°C for 2 hours followed by an overnight incubation at 65°C. DNA was purified using a Qiagen PCR cleanup kit (Qiagen 28104) and subsequently subject to RT-qPCR as described above. Fold-change was calculated using the ΔΔCT method as described above. The ΔCT value for each samples was based off of the CT value for the *trpC* promoter, which has not been shown to be regulated by *whiB7,* nor is it in close proximity to a *whiB7*-regulated gene. The ΔΔCT was then calculated by comparing each experimental sample (i.e. *aspC* knockdown) to the relevant control strain (i.e. non-targeting CRISPRi). Samples derived from a merodiploid *M. smegmatis* strain expressing an untagged *whiB7* did not show any appreciable signal above background.

### Data availability

Raw sequencing data are deposited to the NCBI Short Read Archive under project numbers PRJNA970266 and PRJNA970343.

## SUPPLEMENTAL MATERIALS

**Supplemental Table 1:**
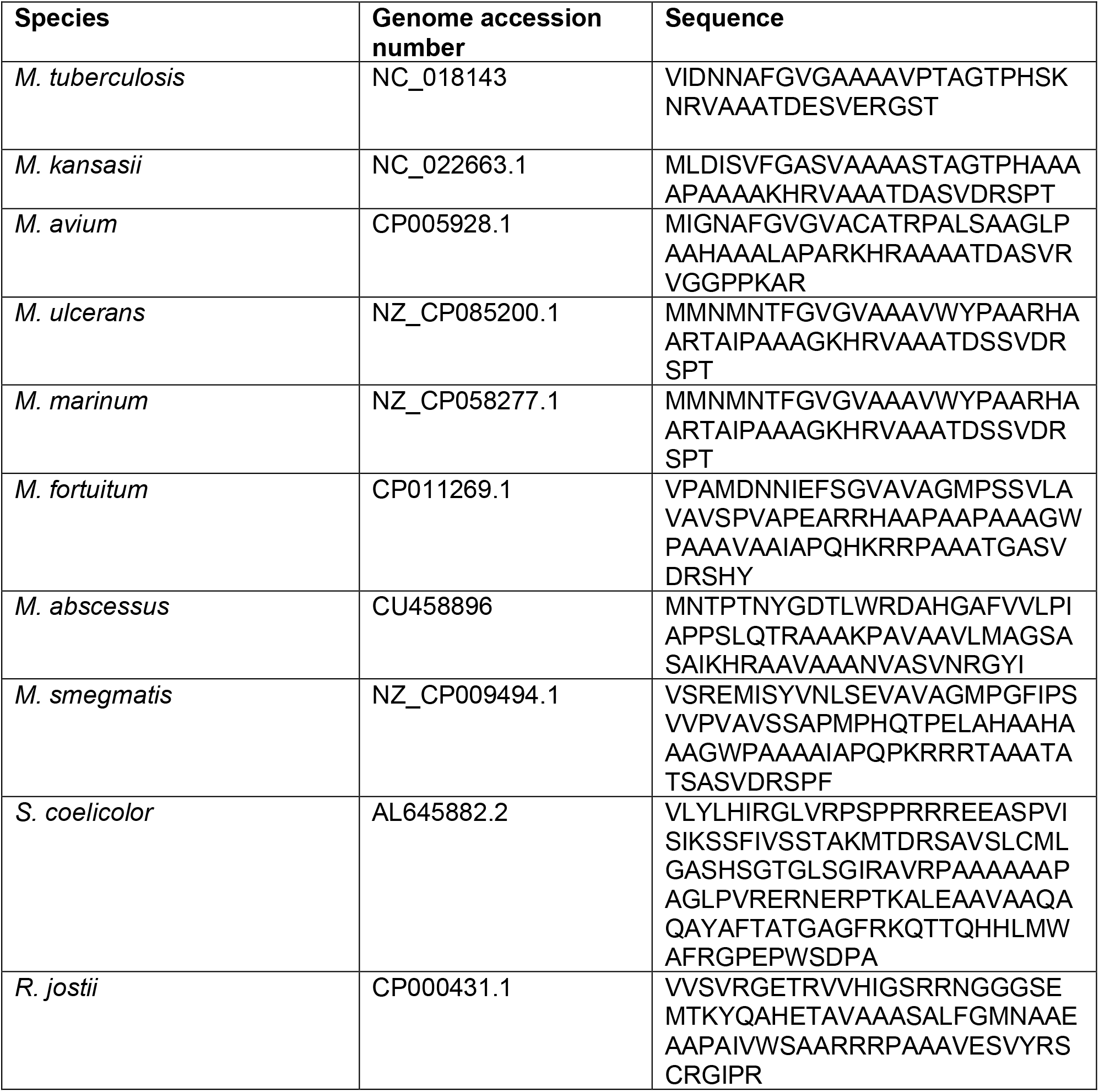
*whiB7* uORF sequences of diverse actinobacteria.

**Supplemental Table 2:** Antimicrobial compounds used in this study.

| **Antibiotic** | **Product number** |
| --- | --- |
| Amikacin | A0365900 (Sigma Aldrich) |
| Anhydrotetracycline | AC233135000 (Fisher Scientific) |
| Azithromycin dihydrate | PZ0007 (Sigma Aldrich) |
| Bedaquiline | A12327-5 (AdooQ Biosciences) |
| Bortezomib | 5043140001 (Sigma Aldrich) |
| Capreomycin Sulfate | PHR1716 (Sigma Aldrich) |
| Chloramphenicol | C-105 (GoldBio) |
| Clarithromycin | C9742 (Sigma Aldrich) |
| Clindamycin hydrochloride | C-175-10 (GoldBio) |
| Ecumicin | NA |
| Ethambutol dihydrochloride | E4630 (Sigma Aldrich) |
| Fusidic acid sodium salt | F0881 (Sigma Aldrich) |
| Isoniazid | I3377 (Sigma Aldrich) |
| Kanamycin | K-120-50 (GoldBio) |
| Levofloxacin | 28266 (Sigma Aldrich) |
| Lincomycin hydrochloride | 62143 (Sigma Aldrich) |
| Linezolid | SML1290 (Sigma Aldrich) |
| PBTZ169 | 22202 (Cayman Chemical) |
| PA-824 (Pretomanid) | SML1290 (Sigma Aldrich) |
| Rifampicin | R0079 (TCI) |
| Spectinomycin dihydrochloride | S-140-5 (GoldBio) |
| Streptomycin sulfate | S-150-100 (GoldBio) |
| Tigecycline hydrate | PZ0021 (Sigma Aldrich) |
| Vancomycin hydrochloride | V2002 (Sigma Aldrich) |
| Zeocin | J67140-8EQ (Alfa Aesar) |
| 6-FABA | 443026 (Sigma Aldrich) |

**Supplemental Figure 1.**
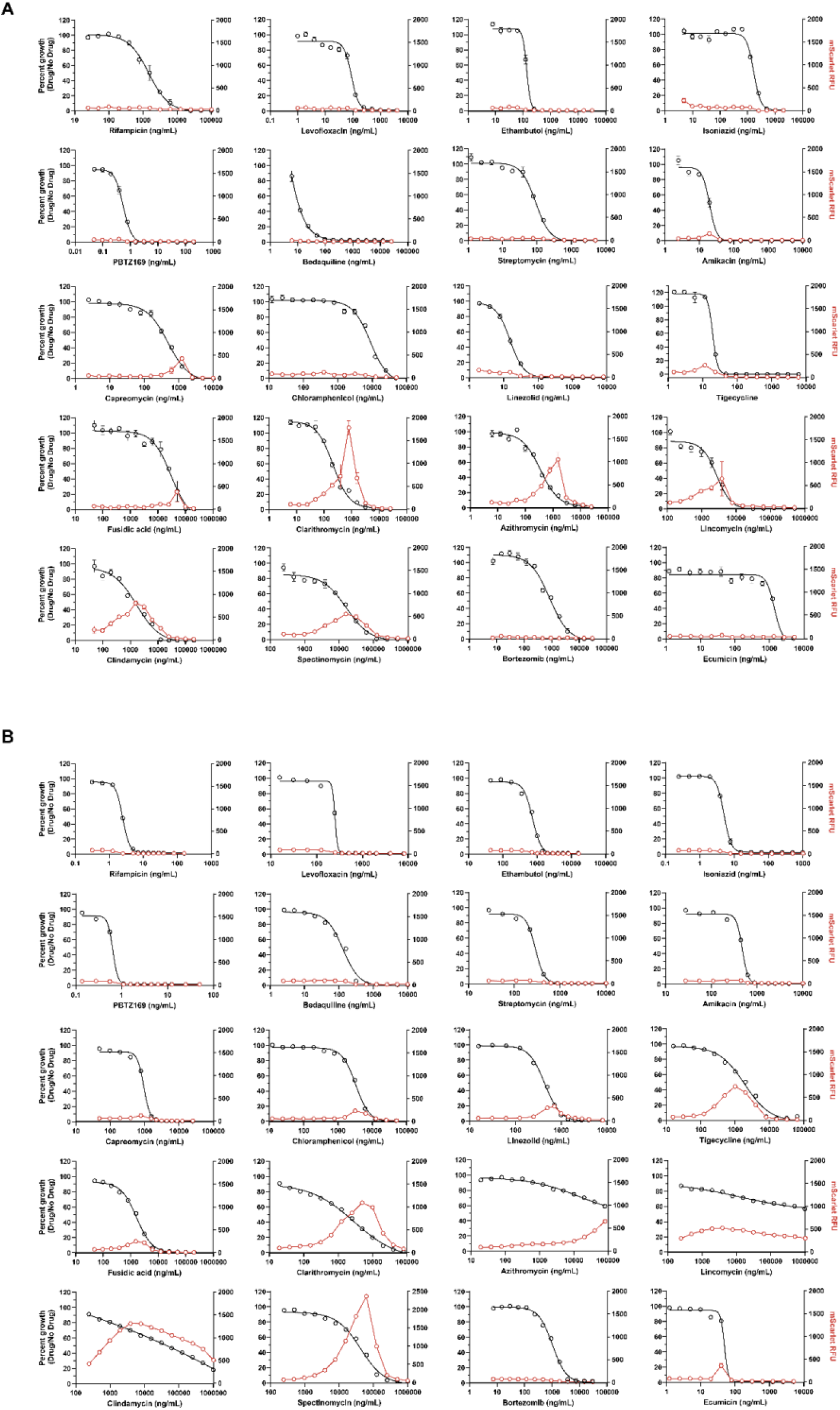
w*h*iB7 activation in response to diverse antibiotics. Dose response curves of the *PwhiB7:mScarlet* reporter strain for the indicated drugs in (A) *M. smegmatis* and (B) *M. tuberculosis*. Drug dose-response curves (percent growth) are shown in black and mScarlet fluorescence (RFU) are shown in red.

**Supplemental Figure 2.**
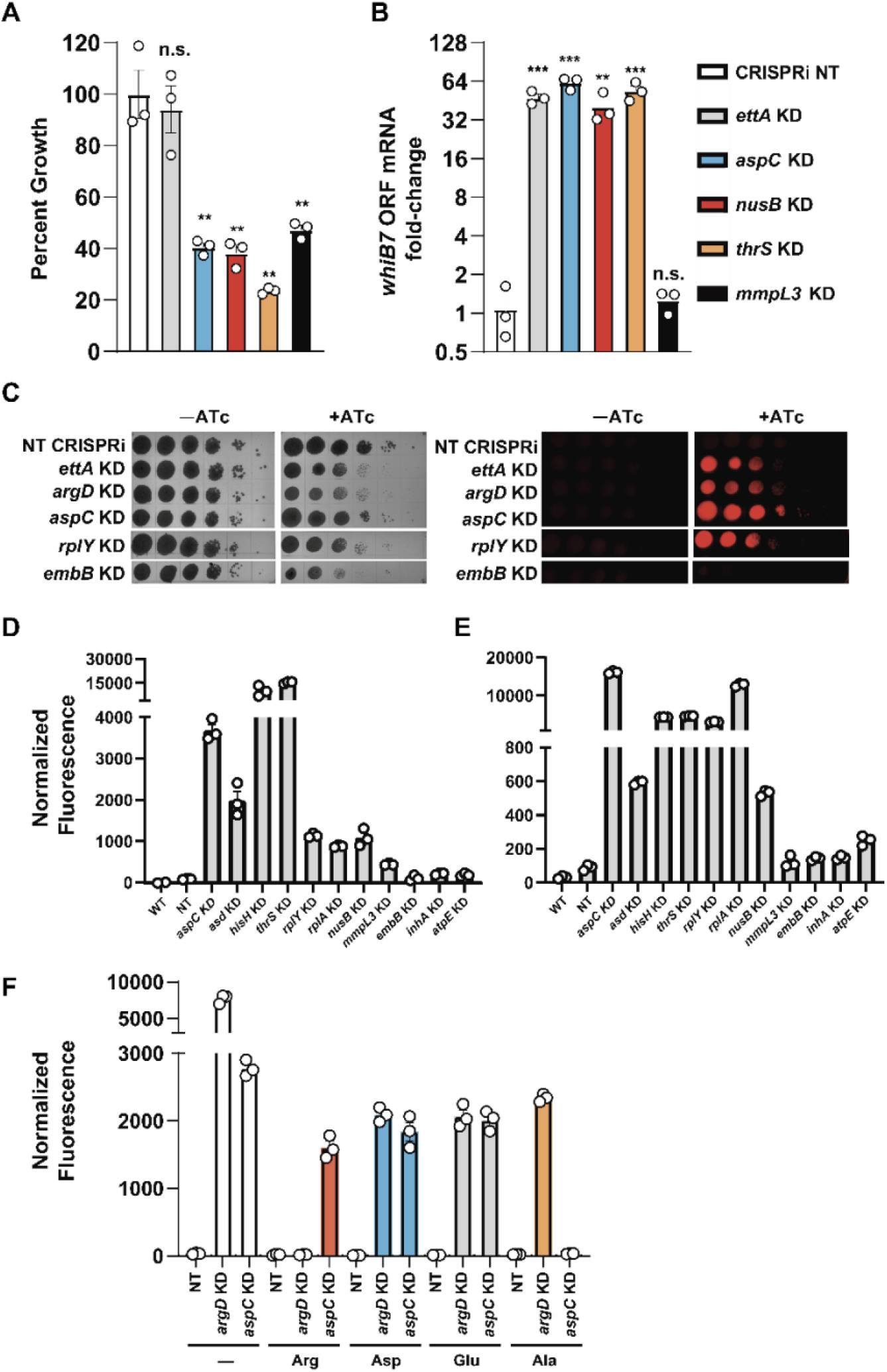
Validation of *whiB7* activation screen. (A,B) Relative growth (A) and *whiB7* ORF mRNA fold-change (B) of the indicated *M. smegmatis* CRISPRi strains 18 hours after addition of ATc (mean ± s.e.m., n = 3 biological replicates). *whiB7* ORF mRNA fold change is relative to *sigA* and normalized to the non-targeting CRISPRi strain. Statistical significance with respect to the non-targeting CRISPRi strain was calculated using a Student’s t-test, **P< 0.01, ***P< 0.001, n.s. = non-significant. (C) Growth and *Pwhib7:mScarlet* induction of the indicated *M. smegmatis* CRISPRi strains monitored by spotting serial dilutions of each strain on the indicated media. Plates were imaged with standard grayscale photography (left) and fluorescent mScarlet imaging (right). (D) Normalized fluorescence of the indicated *M. smegmatis Pwhib7:mScarlet* CRISPRi strains 24 hours after addition of ATc (mean ± s.e.m., n = 3 replicates). (E) Normalized fluorescence of the indicated *M. tuberculosis Pwhib7:mScarlet* CRISPRi strains 8 days after addition of ATc (mean ± s.e.m., n = 3 replicates). (F) Normalized fluorescence of the indicated *M. smegmatis Pwhib7:mScarlet* CRISPRi strains 24 hours after addition of ATc (mean ± s.e.m., n = 3 replicates). The indicated amino acids were added to the culture media at 1 mM.

**Supplemental Figure 3.**
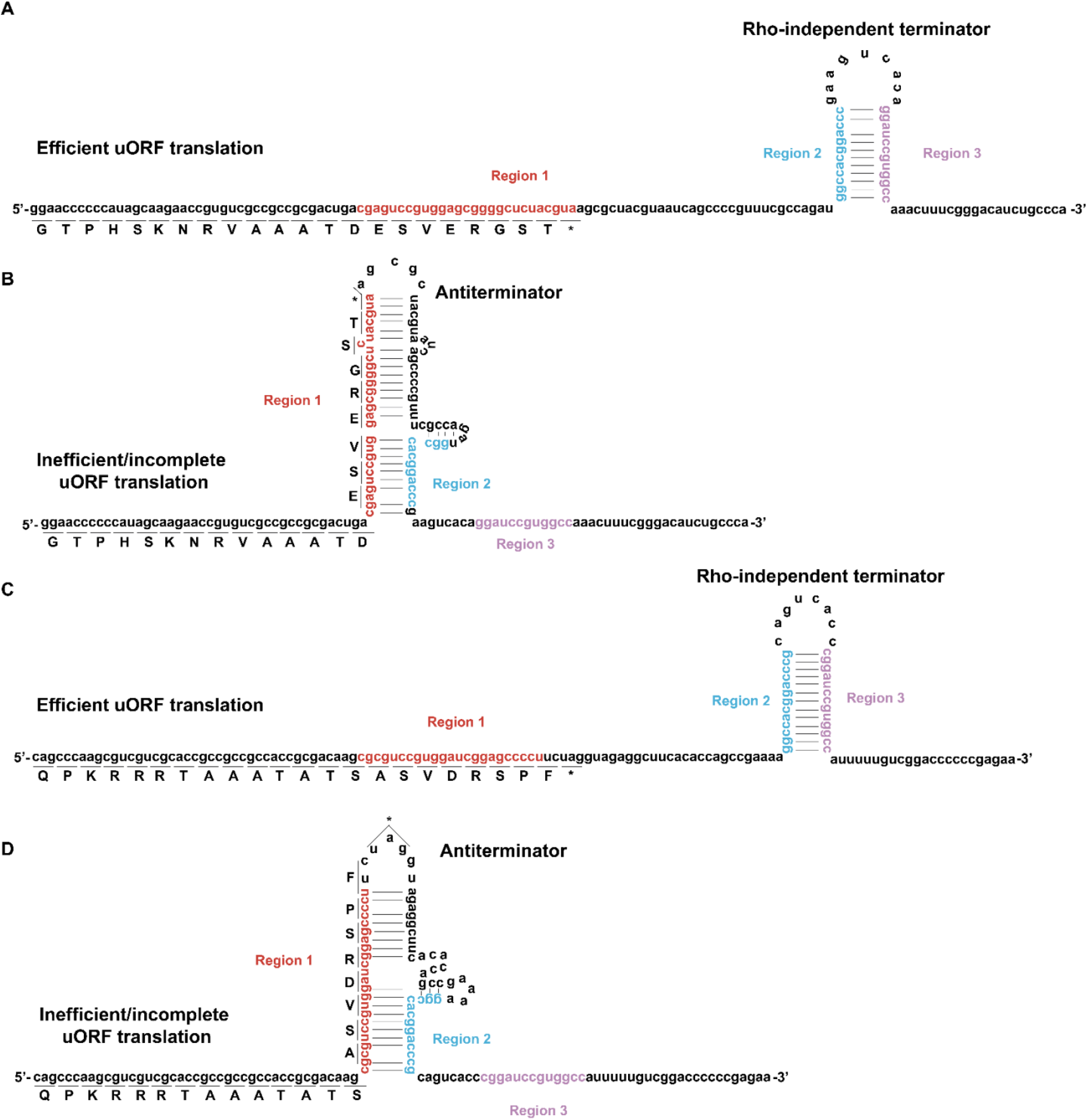
w*h*iB7 5’ regulatory regions of *M. tuberculosis* and *M. smegmatis*. Predicted RNA secondary structures of the *whiB7* 5’ regulatory region during a state of efficient uORF translation (A,C) vs. a state of incomplete or inefficient uORF translation (B,D). Structures for both *M. tuberculosis* (A,B) and *M. smegmatis* (C,D) were predicted based on the work of Lee et al.^22^. For the sake of simplicity, the N-terminal coding sequence of the uORF is not shown. Note that the mutations to produce both the Trp-less and His-less *M. smegmatis* uORF variants occur N-terminal to the sequence depicted in panel (D) and thus should not impact formation of the predicted antiterminator structure.

**Supplemental Figure 4.**
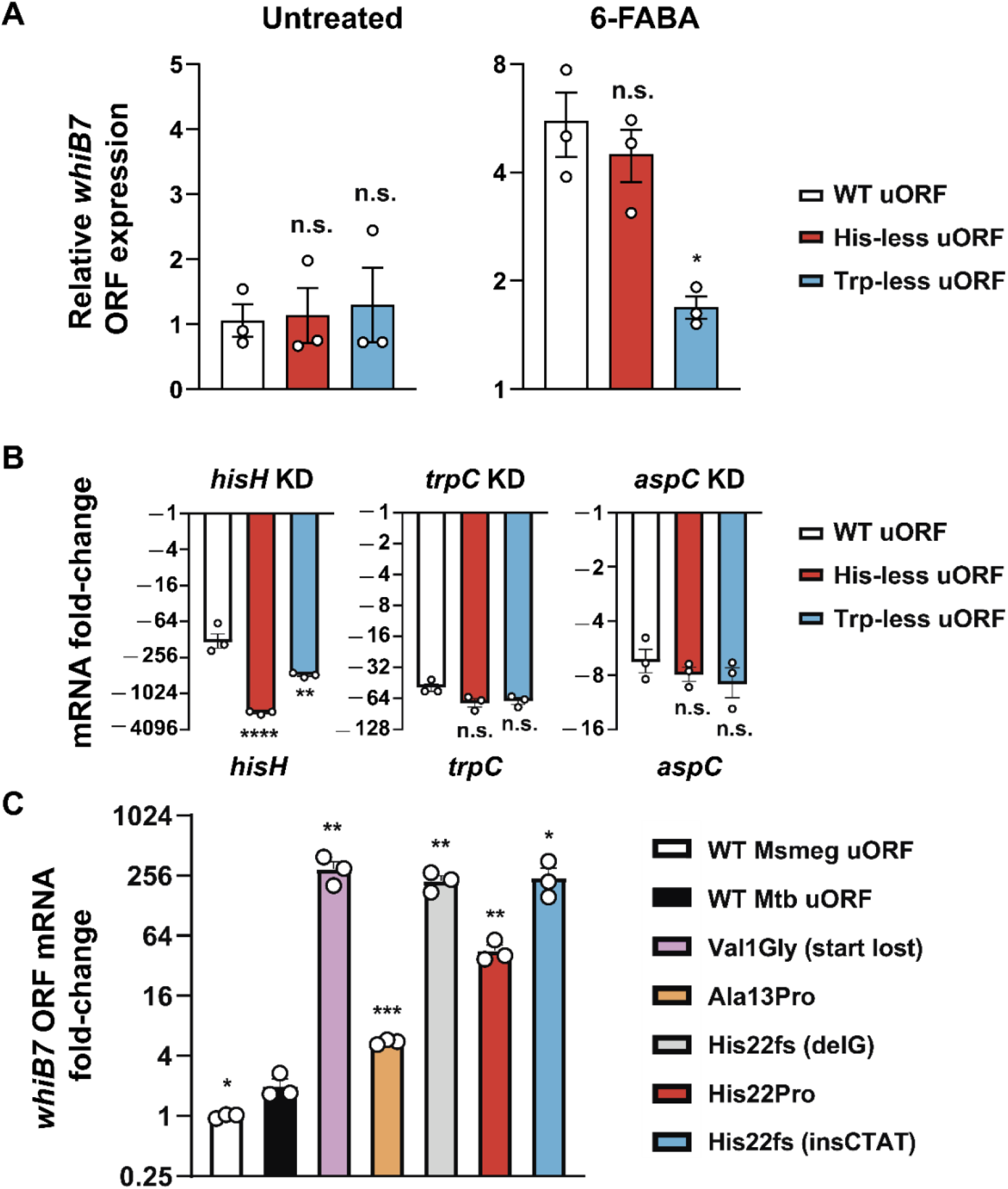
(A) Basal *whiB7* ORF levels are not altered in the *whiB7* uORF variants (Untreated), and deprivation of tryptophan by or 6-FABA treatment (6-FABA) induces *whiB7* expression in the WT and His-less uORF strains but significantly less whiB7 expression in the Trp-less uORF strain. Relative *whiB7* ORF mRNA levels for *M. smegmatis* uORF variants following 15 hours of treatment with 6-FABA (40 ug/mL; mean ± s.e.m., n = 3 biological replicates). *whiB7* ORF mRNA fold change is relative to *sigA* and normalized to the wild-type *whiB7* uORF strain (left panel) or the respective uORF variant without drug treatment (right panel). Statistical significance with respect to the WT *whiB7* uORF strain was calculated using a Student’s t-test; *P< 0.05, n.s. = non-significant. (B) The failure to induce *whiB7* in the uORF variants is not a result of different levels of target gene knockdown (*hisH*, *trpC*, *aspC*). Relative mRNA levels of the indicated genes 15 hours after addition of ATc for the *M. smegmatis* uORF variant CRISPRi strains shown in Figure 4F (mean ± s.e.m., n = 3 biological replicates). mRNA fold-change for the indicated gene at the bottom of each graph was calculated relative to *sigA* and based on the respective non-targeting CRISPRi strain for each uORF variant. Statistical significance with respect to the WT *whiB7* uORF strain was calculated for each knockdown strain using a Student’s t-test; **P< 0.01, ****P< 0.0001, n.s. = non-significant. (C) Identification of rare *whiB7* activating mutations in the uORF of clinical Mtb isolates. Relative *whiB7* ORF mRNA levels for each *M. tuberculosis* uORF variant expressed in the Δ*whiB7 M. smegmatis* strain (mean ± s.e.m., n = 3 biological replicates). mRNA was harvested from log-phase cultures. mRNA fold-change is relative to *sigA* and calculated in comparison to the wild-type *M. smegmatis* uORF strain. Statistical significance with respect to the WT *M. tuberculosis* uORF strain was calculated using a Student’s t-test; *P<0.05, **P< 0.01, ***P< 0.001.

**Supplemental Figure 5.**
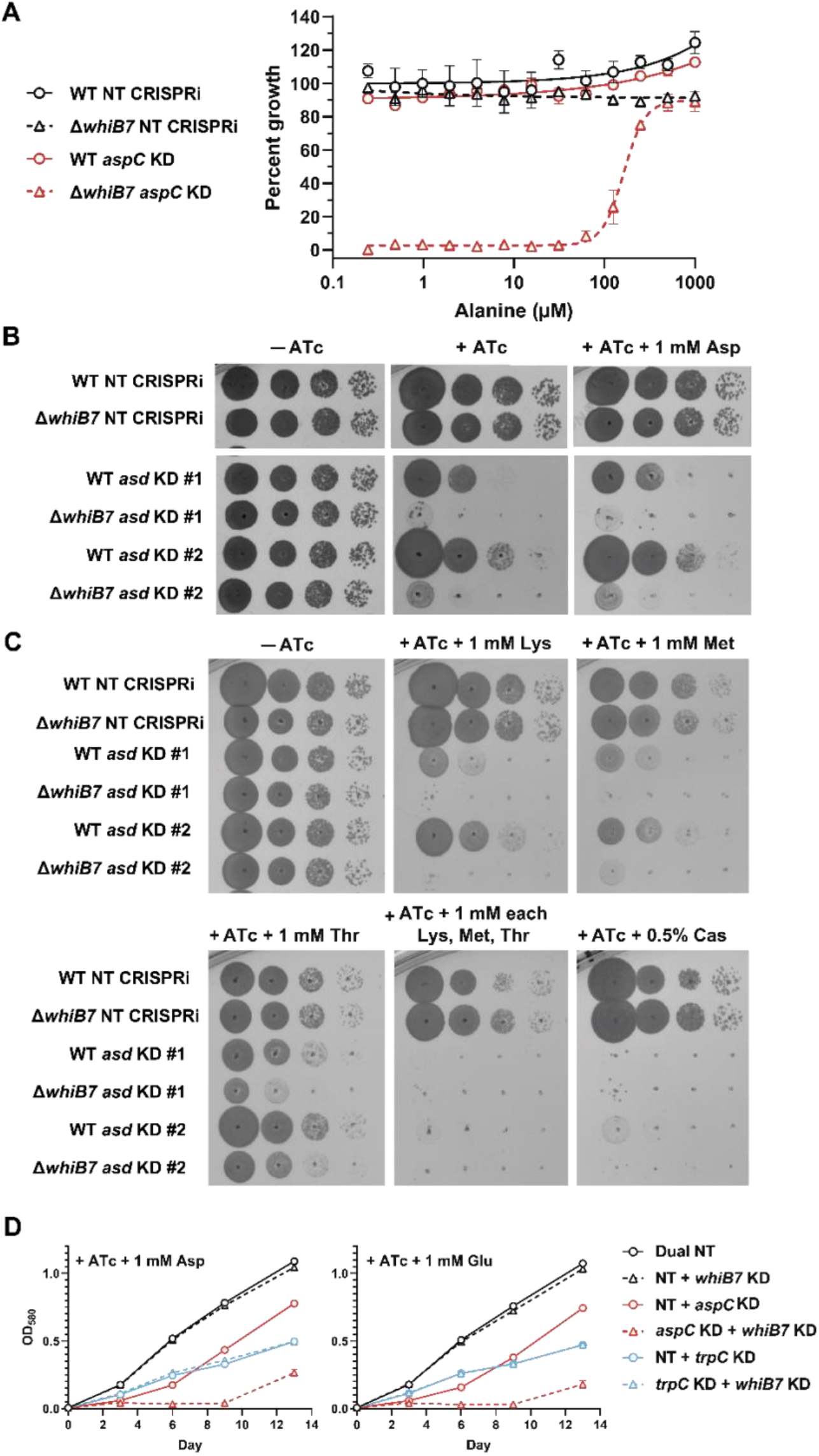
a*s*pC and *asd* are more sensitive to CRISPRi inhibition in Δ*whiB7*. (A) Alanine dose response curves (mean ± s.e.m., n = 3 replicates) for *M. smegmatis aspC* CRISPRi strains in both the wild-type (black) or Δ*whiB7* (red) background. (B-C) Growth of the indicated *M. smegmatis* CRISPRi strains monitored by spotting serial dilutions of each strain on the indicated media. Supplemental amino acids were added at 1 mM, except casamino acids (Cas), which were added at 0.5% w/v. (D) Growth of the indicated *M. tuberculosis* strains in 7H9 + ATc media supplemented with either aspartate or glutamate (both at 1 mM). Dual NT represents a CRISPRi plasmid encoding two non-targeting sgRNAs; NT + *whiB7* KD represents a CRISPRi plasmid encoding a single non-targeting sgRNA and a *whiB7* targeting sgRNA; *aspC* KD + *whiB7* KD represents a CRISPRi plasmid encoding one sgRNA targeting *aspC* and a separate sgRNA targeting *whiB7*.

